# Myocardin-related transcription factors regulate morphogenetic events in vertebrate embryos by controlling F-actin organization and apical constriction

**DOI:** 10.1101/2023.09.27.559818

**Authors:** Keiji Itoh, Olga Ossipova, Miho Matsuda, Sergei Y. Sokol

## Abstract

Myocardin-related transcription factors (Mrtfa and Mrtfb), also known as megakaryoblastic leukemia proteins (Mkl1/MAL and Mkl2), associate with serum response factor (Srf) to regulate transcription in response to actin dynamics, however, the functions of Mrtfs in early vertebrate embryos remain largely unknown. Here we document the requirement of Mrtfs for blastopore closure at gastrulation and neural plate folding in Xenopus early embryos. Both stimulation and inhibition of Mrtf activity caused similar gross morphological phenotypes, yet the effects on F-actin distribution and cell behavior were different. Suppressing Mrtf-dependent transcription reduced overall F-actin levels and inhibited apical constriction during gastrulation and neurulation. By contrast, constitutively active Mrtf caused tricellular junction remodeling and induced apical constriction in superficial ectoderm. The underlying mechanism appeared distinct from the one utilized by known apical constriction inducers. We propose that the regulation of apical constriction is among the primary cellular responses to Mrtf. Our findings highlight a dedicated role of specific transcription factors, Mrtfs, in early morphogenetic processes.

## Introduction

Early vertebrate development involves the specification of embryonic cell fates and the regulation of morphogenetic processes that place cells and tissues into their prospective positions. Actin remodeling plays a major role in the control of cell shape and cell motility (Buracco et al., 2019; Pollard and Cooper, 2009). In addition, actin dynamics regulates gene transcription (Olson and Nordheim, 2010) that contributes to the interplay between cell fate specification and the cell movements accompanying morphogenesis. Early vertebrate embryos serve as an excellent *in vivo* model for studying these mechanisms, because they exhibit diverse cell behaviors, such as epiboly, apical constriction, tissue involution and convergent extension, that are needed for gastrulation and neural tube closure (Keller et al., 2003; Keller and Sutherland, 2020; Vijayraghavan and Davidson, 2017).

The myocardin-related transcription factors Mrtfa and Mrtfb, also known as megakaryoblastic leukemia 1 and 2 (Mkl1 and Mkl2), respectively, are conserved regulators of cellular actin dynamics, linking actin polymerization and transcriptional control. Accumulating evidence points to Mrtfs as having important functions in normal development and disease. In mice, Mrtfs are necessary for development of smooth muscle in the heart (Cenik et al., 2016; Li et al., 2005; Mokalled et al., 2015)*. Mrtfa^-/-^* mice are viable (Li et al., 2006; Sun et al., 2006b) whereas *Mrtfb^-/-^*mice die between E13.5-E15.5 due to cardiovascular defects (Li et al., 2005; Oh et al., 2005). In humans, Mrtfa is primarily associated with acute megakaryoblastic leukemia in children (Mercher et al., 2001).

Mrtfs act as transcriptional cofactors by forming a complex with Serum response factor (Srf), an evolutionary conserved protein (Cen et al., 2003; Miralles et al., 2003; Olson and Nordheim, 2010; Wang et al., 2002). Mrtfs contain a transcriptional activation domain, whereas Srf has a conserved DNA-binding domain. The binding of the Mrtf/Srf complex to CArG motifs at target gene promoters leads to transcriptional activation. Not surprisingly, *Srf^-/-^* mice die due to gastrulation and mesoderm specification defects (Arsenian et al., 1998; Niu et al., 2005). Conditional Srf knockout mice exhibit abnormal angiogenesis, neural, cardiac and craniofacial development (Franco et al., 2008; Niu et al., 2005; Stritt et al., 2009; Vasudevan and Soriano, 2014). The formation of the active Srf/Mrtf transcriptional complex in the nucleus is inhibited by monomeric G-actin that sequesters Mrtfs in the cytoplasm (Cen et al., 2003; Miralles et al., 2003). The removal of the G-actin-binding domain can produce a constitutively active form of Mrtf (Geneste et al., 2002; Miralles et al., 2003; Weissbach et al., 2016). Consistent with this mechanism, F-actin polymerization reduces G-actin levels and promotes Mrtf nuclear localization and transcriptional activation of target genes (Esnault et al., 2014; Medjkane et al., 2009). Supporting this model, Mrtf and Srf regulate motility of various cell lines (Kim et al., 2017; Liao et al., 2014; Medjkane et al., 2009; Morita et al., 2007; Seifert and Posern, 2017) and border cell migration in *Drosophila* (Salvany et al., 2014; Somogyi and Rorth, 2004).

In this study, we investigate Mrtf functions during *Xenopus* embryonic development. In agreement with previous studies, we find that putative Mrtf target genes are largely associated with actin cytoskeleton and cell junction remodeling. Consistent with the proposed roles of Mrtfs in controlling cell shape and behavior, we show that Mrtfs modulate blastopore closure during gastrulation and neural plate folding. Constitutively active and dominant interfering Mrtf constructs had opposite effects on *in vivo* Srf reporter activity, however, both active and inhibitory constructs caused gastrulation and neurulation defects. We find that a major cellular behavior affected by Mrtf during morphogenesis is apical construction. The active form of Mrtf triggered ectopic apical constriction in superficial ectoderm, likely by affecting F-actin organization at tricellular junctions. Our observations establish an essential role of Mrtfs in controlling cell shape during vertebrate gastrulation and neural tube closure.

## RESULTS

### Myocardin-related transcription factors modulate Srf-dependent transcription in early *Xenopus* embryos

Mrtf proteins contain multiple functional domains (**Fig. 1A**)(Miralles et al., 2003; Miranda et al., 2021; Zaromytidou et al., 2006). The binding of Mrtf by G-actin is mediated by the N-terminal RPEL motifs (Miralles et al., 2003; Vartiainen et al., 2007), while Srf associates with the basic (B) region (Cen et al., 2003; Miralles et al., 2003; Zaromytidou et al., 2006). The SAP (<u>S</u>AF-A/B, <u>A</u>cinus and <u>P</u>ias) domain has been implicated in the interaction with DNA (Aravind and Koonin, 2000; Kipp et al., 2000), whereas the conserved leucine-zipper motif mediates dimerization. Importantly, Mrtfs contain a transcriptional activation domain at the C-terminus (Cen et al., 2003; Miralles et al., 2003; Zaromytidou et al., 2006).

**Fig. 1.**
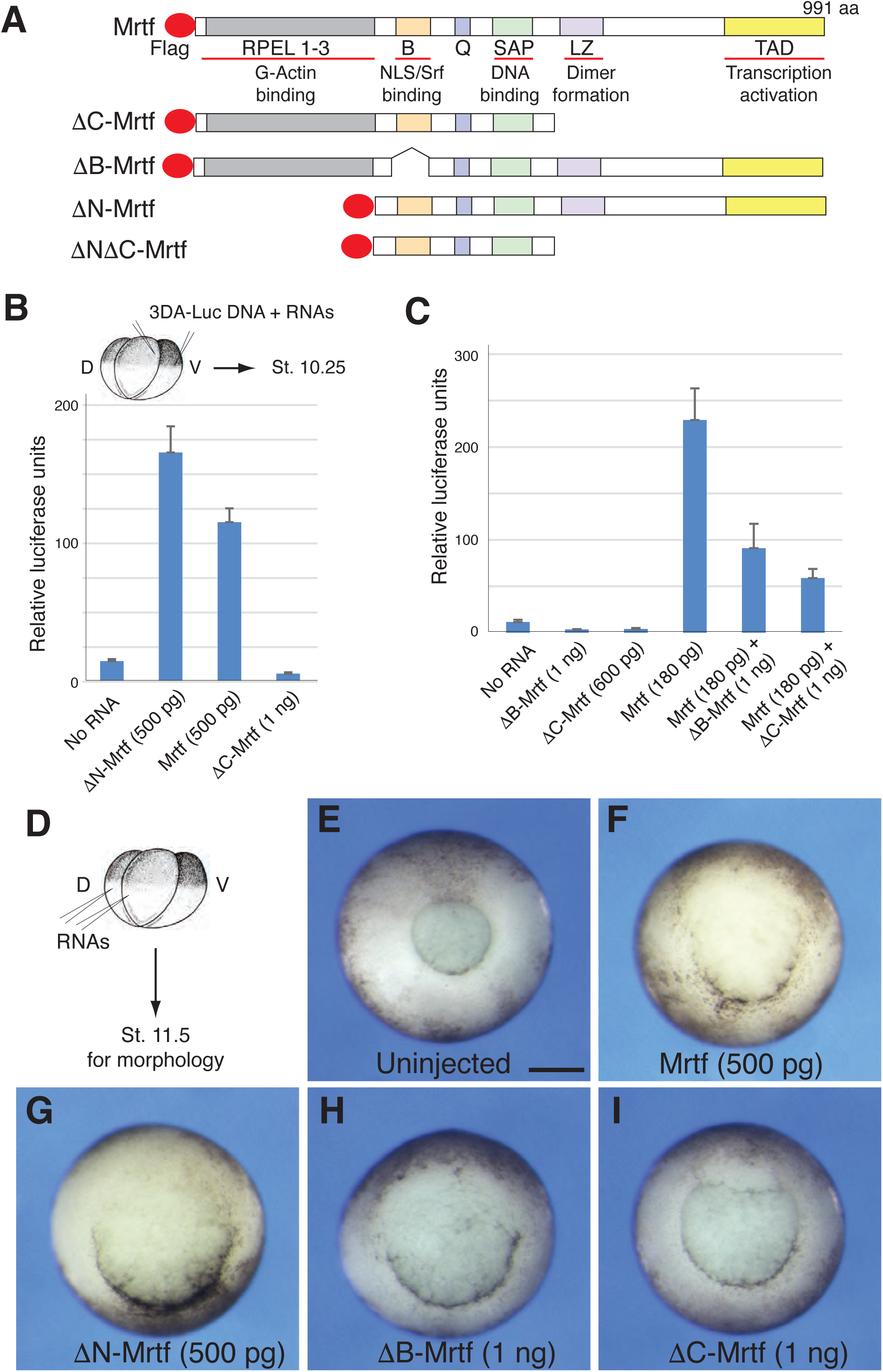
Distinct effects of Mrtf constructs on Srf reporter activation and blastopore closure during gastrulation. A, Mrtfa constructs used in this study: B, Basic region that contains nuclear localization sequence and binds Srf. Q, Glutamine-rich region. SAP, DNA binding domain. LZ, leucine-zipper dimerization domain. TAD, transcriptional activation domain. B, C, Experimental scheme for transient *3DA-Luc* reporter assays in *Xenopus* ectoderm. B, Transcriptional activation of *3DA-Luc* DNA by different Mrtf constructs. C, ΔB-Mrtf and ΔC-Mrtf inhibit endogenous and exogenous Mrtf activity. Means +/- s.d. are shown. D-I. Dorsal blastopore formation is inhibited by both constitutively active and dominant interfering Mrtf constructs. D, Experimental scheme. Two dorso-vegetal sites of four-cell embryos were injected with the indicated RNAs. E, Uninjected control embryo, vegetal view. (F-I) Dorsal blastopore defects in representative embryos expressing Mrtf (F), ΔN-Mrtf (G), ΔB-Mrtf (H) and ΔC-Mrtf (I). Vegetal view is shown, dorsal is up, RNA doses indicated. Images are representative of 3-4 different experiments. Phenotypes were scored when normal sibling embryos reached stage 11.5. Scale bar: 300 μm.

To assess whether the mechanism of Mrtf signaling that was established in tissue culture cells is preserved in vertebrate embryos, we generated several mutated variants of *Xenopus* Mrtfa lacking specific regulatory domains and will refer to them for simplicity as Mrtf constructs (**Fig. 1A**). RNAs encoding various Mrtf constructs have been injected into *Xenopus* early embryos to examine their effects on Srf transcriptional activation. For this assay, we used *p3DA-Luc*, a luciferase reporter containing three Srf binding sites, which is specifically activated by the Mrtf/Srf complex (Busche et al., 2008; Hill et al., 1995).

Consistent with the inhibitory role of G-actin on Mrtf, ΔN-Mrtf lacking the actin-binding RPEL motifs activated *p3DA-Luc* more strongly than wild-type Mrtf did (**Fig. 1B**). The Srf reporter was not activated by ΔC-Mrtf (lacking the leucine zipper and transcriptional activation domain) and ΔB-Mrtf deficient in Srf binding **(Fig. 1C**). Additionally, the two constructs suppressed reporter activation by wild-type Mrtf **(Fig. 1C**), exhibiting a dominant interfering activity. A control reporter lacking Srf-binding CArG sequences was not stimulated by any of the Mrtf constructs (data not shown).

To identify putative Mrtf gene targets. We carried out RNA sequencing of ectoderm cells expressing ΔN-Mrtf. This analysis revealed that the top Mrtf-induced genes encode various actins and actin-associated proteins (**Fig. S1A, B**). Confirming the results of the RNA sequencing, several of the targets, including *acta2, actc1* and *myl3,* were validated by quantitative RT-PCR (**Fig. S1C**). These findings indicate that *in vivo* Mrtf targets in *Xenopus* early embryos closely match the gene targets previously identified *in vitro* in mammalian cultured cells (Esnault et al., 2014; Medjkane et al., 2009; Selvaraj and Prywes, 2004; Sun et al., 2006a).

These experiments validated the transcriptional activity of the Mrtf/Srf complex in *Xenopus* embryos and pointed to actins as key cellular targets that underlie Mrtf-dependent morphogenetic processes.

### Regulation of early morphogenetic events by Mrtfs

We next examined the effects of the Mrtf constructs on *Xenopus* gastrulation. Mrtf mRNAs were injected into dorsovegetal blastomeres of 4-8-cell embryos targeting the cell population that forms the dorsal blastopore (**Fig. 1D-I**). During normal gastrulation, blastopore first appears at the dorsal marginal zone, then propagates laterally and ventrally and closes at the end of gastrulation (Keller, 1991). Both putative dominant interfering Mrtf constructs, ΔC-Mrtf and ΔB-Mrtf, inhibited dorsal blastopore formation (**Fig. 1H, I)**. Of note, the full-length and constitutively active ΔN-Mrtf also interfered with blastopore appearance and closure (**Fig. 1F, G)**. At later stages, both active and inhibitory forms of Mrtf caused neural tube closure defects and head abnormalities (**Fig. S2**). The tested Mrtf constructs were expressed similarly, with the exception of ΔN-Mrtf that had a lower level of expression (**Fig. S3**). Similar observations have been made using derivatives of human Mkl1, kindly provided by Ron Prywes (Columbia University) (data not shown). Together, our findings indicate that the proper levels of Mrtf activity are critical for gastrulation and later morphogenesis.

To confirm a role of Mrtfs in early morphogenetic events using a different approach, we designed splicing-blocking MOs that are specific for *mrtfa* or *mrtfb* genes. In both cases, the successful targeting of the junction sequences between the second intron and the third exon has been validated by RT-PCR (**Fig. 2A-C**). Injections of Mrtfa MO, Mrtfb MO or both MOs caused defects in blastopore closure (**Fig. 2D-H**). In addition, at later stages, we observed neural tube defects (**Figs. 2I-M**). Though both MOs showed similar phenotypes at gastrulation, Mrtfa MO showed stronger neural tube defects. These observations establish critical roles of Mrtfs in morphogenetic processes that accompany gastrulation and neural tube closure.

**Fig. 2.**
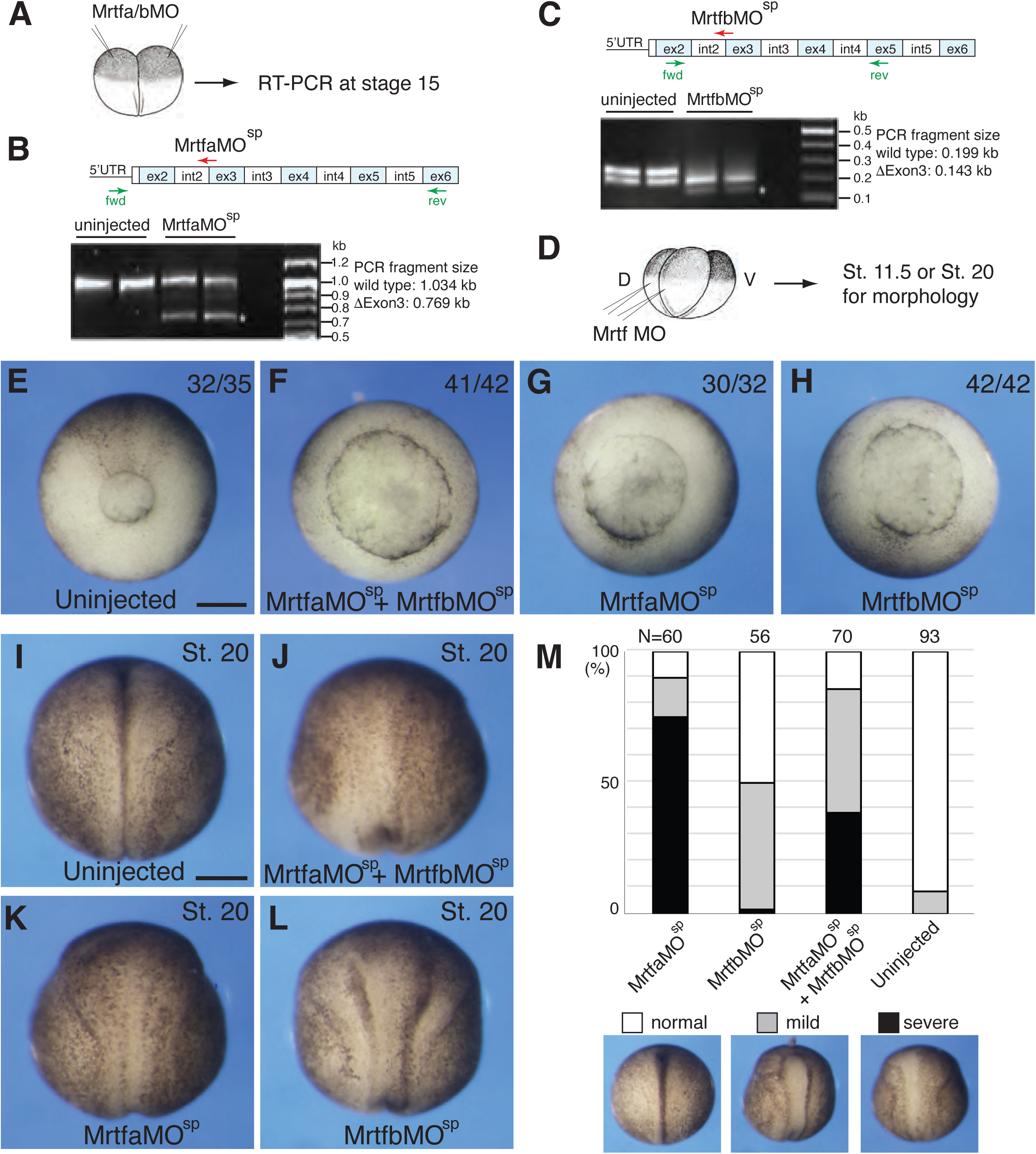
Mrtf is required for Xenopus gastrulation and neurulation. A-C, Validation of *Mrtf* knockdown. A, Experimental scheme. Embryos were injected animally with Mrtfa-MO^sp^ or Mrtfb-MO^sp^ (25 ng each). B, C. Mrtfa or Mrtfb gene structure with indicated exon (ex) and intron (int) boundaries is shown above each gel. Both Mrtfa-MO^sp^ and Mrtfb-MO^sp^ target the junction of intron 2 and exon 3. RT-PCR with specific primers show the removal of exon 3 in the morphants. Each group was analyzed in duplicate samples, control uninjected embryos are on the left. The results are representative of two different experiments. B, Mrtfa MO^sp^; C, Mrtfb MO^sp^. D-H. Gastrulation defects in Mrtf morphants. Two dorso-vegetal sites of four-cell embryos were injected with Mrtfa-MO^sp^ or Mrtfb-MO^sp^ separately (60 ng each) or in combination (30 ng each) as indicated. E-H, Blastopore formation was assessed when control embryos reached stage 11.5. Vegetal view of representative embryos is shown, dorsal is at the top. The images shown are representative of three different experiments. Phenotype frequencies and the total numbers of scored embryos are indicated for each group. I-M, Morphant phenotypes at neurulation. When control sibling embryos reached stage 20, neural tube closure defects were imaged. I-L, Dorsal view of representative embryos is shown, anterior is at the top. M, Quantification of data shown in I-L. Neural tube closure defects were scored as severe, mild or normal. Data are representative of two to three independent experiments. Scale bars for E-H or J-L: 300 μm.

### Mrtf activity controls F-actin remodeling and cell shape

Given the association of Mrtfs with actin transcription, we hypothesized that Mrtfs may regulate F-actin levels or localization. We therefore assessed the distribution of F-actin in early gastrulae (**Fig. 3**). Consistent with our hypothesis, ΔC-Mrtf reduced apical F-actin levels, consistent with *actin* being a major transcriptional target for Mrtfs (Esnault et al., 2014; Salvany et al., 2014)(**Fig. 3B-B”’**). By contrast, ΔN-Mrtf selectively disrupted the accumulation of F-actin at the tricellular (TCJ) but not bicellular junctions (BCJ) (**Fig. 3C-C”’**). We also found that ΔN-Mrtf reduced the enrichment of Tricellulin, an obligatory TCJ component (Higashi and Miller, 2017), at TCJ (**Fig. S4A-C**). Both Tricellulin and F-actin signals have spread to BCJ, consistent with junctional remodeling (**Fig. S4C**).

**Fig. 3.**
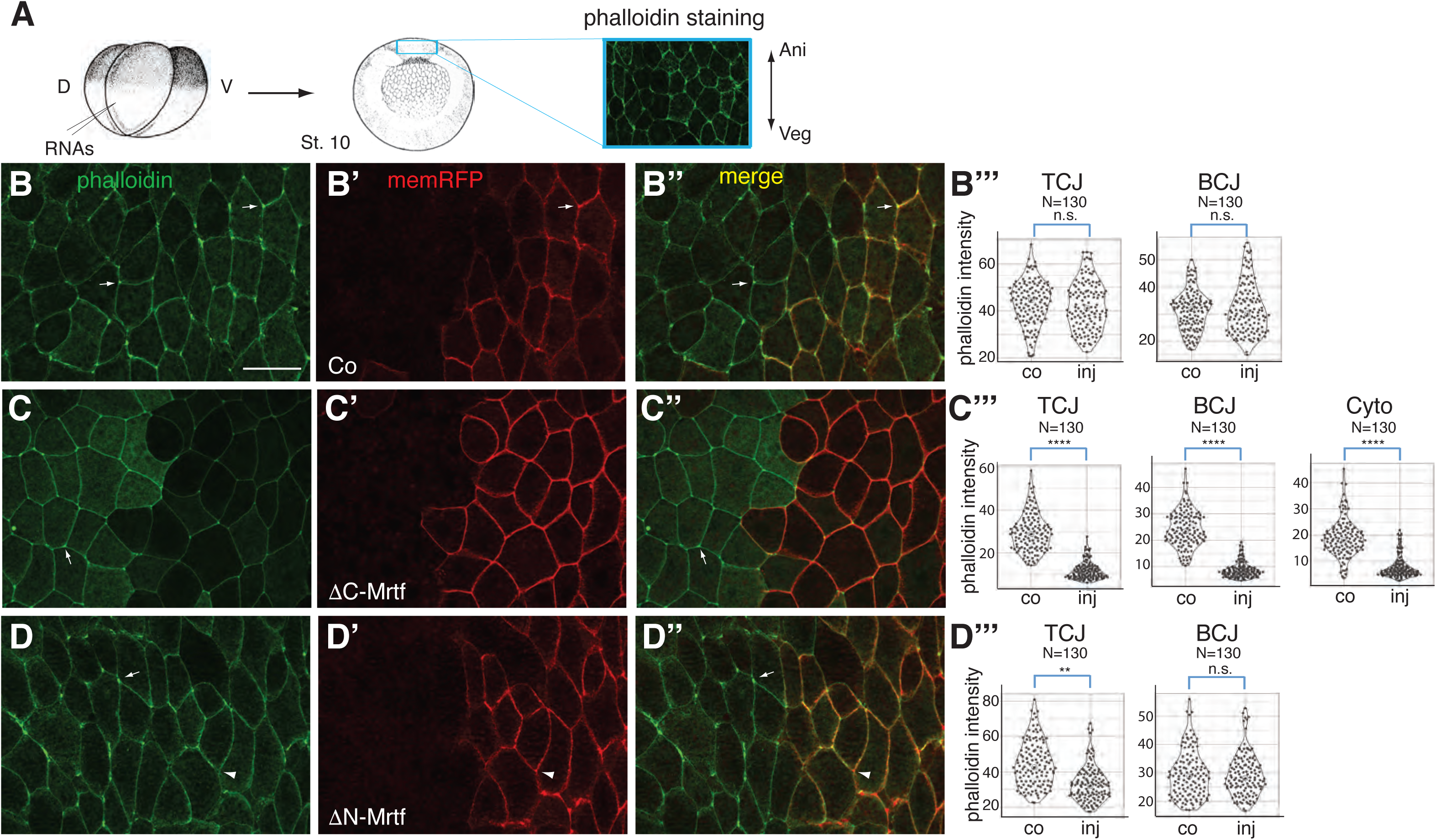
Modulation of Mrtf activity alters F-actin levels and localization. **A.** Experimental scheme is shown on top left. Four-cell embryos were injected into one dorso-vegetal blastomere with membrane RFP (memRFP) RNA (100 pg) alone (B-B”’) or with 1 ng ΔC-Mrtf RNA (C-C”’) or 250 pg ΔN-Mrtf RNA (D-D”’). Dorsal marginal zone (DMZ) explants of the injected embryos at stage 10 were stained with phalloidin. The rectangular area indicated above the dorsal blastopore was imaged. Images of phalloidin (B, C, D), memRFP (B’, C’, D’) or merge (B”, C”, D”) are shown. Phalloidin intensities at tricellular junctions (TCJ), bicellular junctions (BCJ) and cytoplasm (Cyto) were assessed on the control (co) vs the injected (inj) side (B”’, C”’, D”’). Arrows in B-D point to TCJ. Arrowheads in D-D” indicate a TCJ with reduced F-actin. Statistical significance is indicated with Student’s t-test. **p<0.01, ****p<0.0001, n.s. non-significant. Data are representative of 3-4 independent experiments. Scale bar: 50 μm.

Since tricellular junction remodeling may alter cell shape, we carried out live imaging of embryonic ectoderm cells mosaically expressing ΔN-Mrtf and compared them to the mosaic control cells (**Supplementary movies 1, 2**). In stage 11 ectoderm, we consistently observed that ΔN-Mrtf expressing cells had convex shape as compared to the concave shape of control neighboring cells (**Fig. 4A-D**). Notably, apical domain size of ΔN-Mrtf expressing cells was smaller than that of the control cells (**Fig. 4B, C, E**). Our experiments reveal a specific change in cell shape that is consistent with the regulation of the actin cytoskeleton by Mrtf-dependent transcription. The observed changes in cell shape might reflect changes in surface tension or cell adhesion (Maitre et al., 2012; Sedzinski et al., 2016; Winklbauer, 2015), however the clarification of the specific mechanism awaits future studies.

**Fig. 4.**
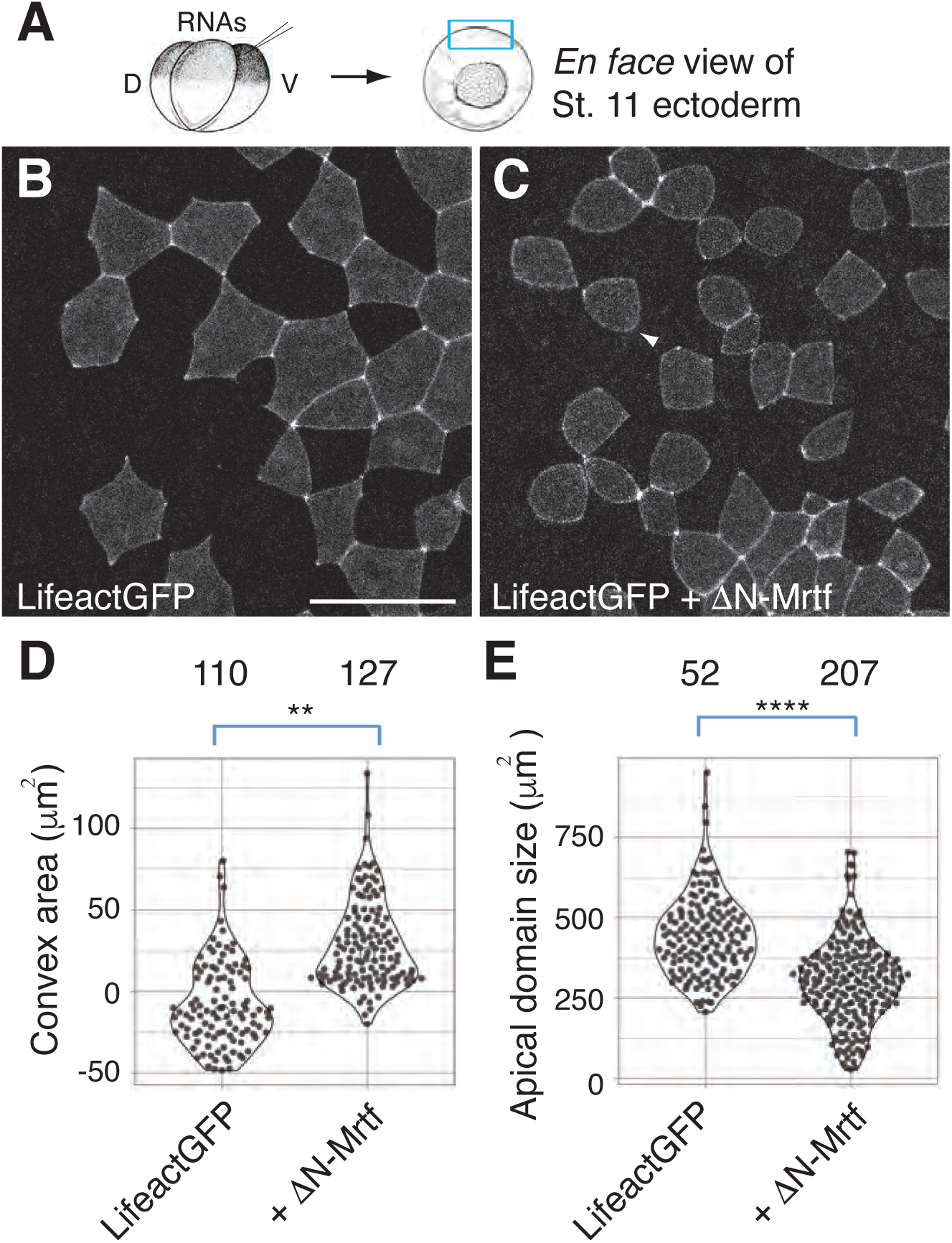
ΔN-Mrtf alters cell shape in the epidermal ectoderm. A. Experimental scheme. Four-cell embryos were injected into one-ventro-animal blastomere with LifeactGFP RNA (100 pg) alone (B) or with 250 pg ΔN-Mrtf RNA (C). Superficial epidermal ectoderm of the injected embryos was imaged at stage 11. Arrowhead in C points to a cell with convex shape. D. Quantification of shape of mosaic GFP expressing cells (the cells contacting no more than two other GFP expressing cells). Convex and concave areas between two junctional spots of GFP were measured. Concave areas of membrane curvature were measured and assigned negative values. Numbers of scored junctions are on the top. **p<0.01. E. Apical domain size of mosaic GFP-expressing cells. Numbers of scored cells are on the top, ****p<0.0001, Student’s t-test. Scale bar: 50 μm.

### Regulation of apical constriction by Mrtf

In our experiments, Mrtf depletion and overexpression of the inhibitory ΔC-Mrtf construct caused defective blastopore closure and delayed neural fold formation (**Figs. 1, 2, Fig. S2**). Notably, both blastopore development and neural groove formation are characterized by the formation of bottle cells, i. e. cells with reduced apical surface. Since we noticed that the apical surface of ΔN-Mrtf-expressing cells is slightly reduced in stage 11 ectoderm (**Fig. 4**), we decided to test the hypothesis that ΔN-Mrtf can directly modulate apical constriction.

We examined more closely how ΔN-Mrtf affects apical constriction in ectoderm cells at later developmental stages after mosaic expression. At stage 12.5-13, ΔN-Mrtf-expressing cells exhibited much smaller apical domain than ΔC-Mrtf-expressing cells (**Fig. 5A-D”, F**). To further demonstrate that ΔN-Mrtf induces apical constriction, time lapse imaging was carried out during the 3.5 hr period between stages 10.5 and 12.5 (data not shown) and apical constriction was observed in 46% of ΔN-Mrtf-expressing cells (n=143). By contrast, less than 4% control cells expressing MyrGFP alone or with ΔC-Mrtf were constricting (n=120 and n=132, respectively) (**Fig. 5G**). The conclusion that that active Mrtf induces ectopic apical constriction has been confirmed in cryosections (**Fig. 6A-D**).

**Fig. 5.**
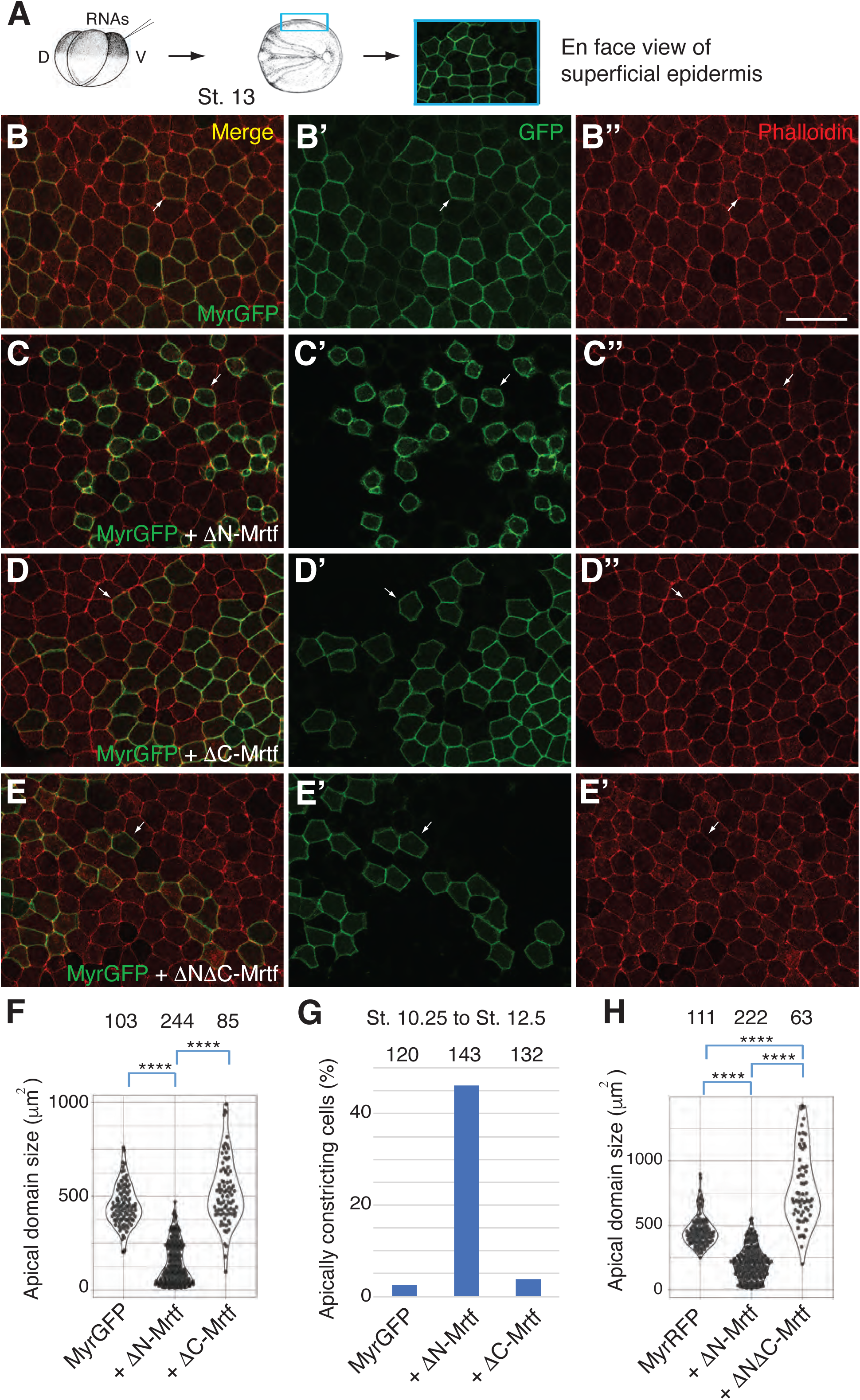
Constitutively active Mrtf induces ectopic apical constriction. A. Experimental scheme. Four-cell embryos were injected into one ventral-animal blastomere with MyrGFP RNA (50 pg) alone (B) or with 250 pg ΔN-Mrtf RNA (C), ΔC-Mrtf RNA (D) or ΔNΔC-Mrtf RNA (E). Superficial epidermal ectoderm of the injected embryos at stage 13 was imaged. Arrows point to the cells mosaically expressing GFP. Scale bar: 50 μm. F. Apical domain size of mosaic cells was measured in a separate experiment with 1 ng ΔC-Mrtf RNA. ****p<0.0001, Student’s t-test. G. Live imaging of superficial ectodermal cells expressing MyrGFP RNA (50 pg) alone or with 250 pg ΔN-Mrtf or ΔC-Mrtf RNA was performed between stage 10.25 and 12.5. Ratios of apical domain size measured at stage 12 to that measured at stage 10.25 were calculated for individual cells. Apically constricting cells were scored as the ones with the ratio of 0.8 or below. Numbers of scored cells are on top of F-H. H. Four-cell embryos were injected with MyrRFP RNA (50 pg) alone or with 250 pg ΔN-Mrtf RNA or 1 ng ΔNΔC-Mrtf RNA as shown in A. Apical domain size of mosaic, RFP expressing cells. ****p<0.0001, Student’s t-test.

**Fig. 6.**
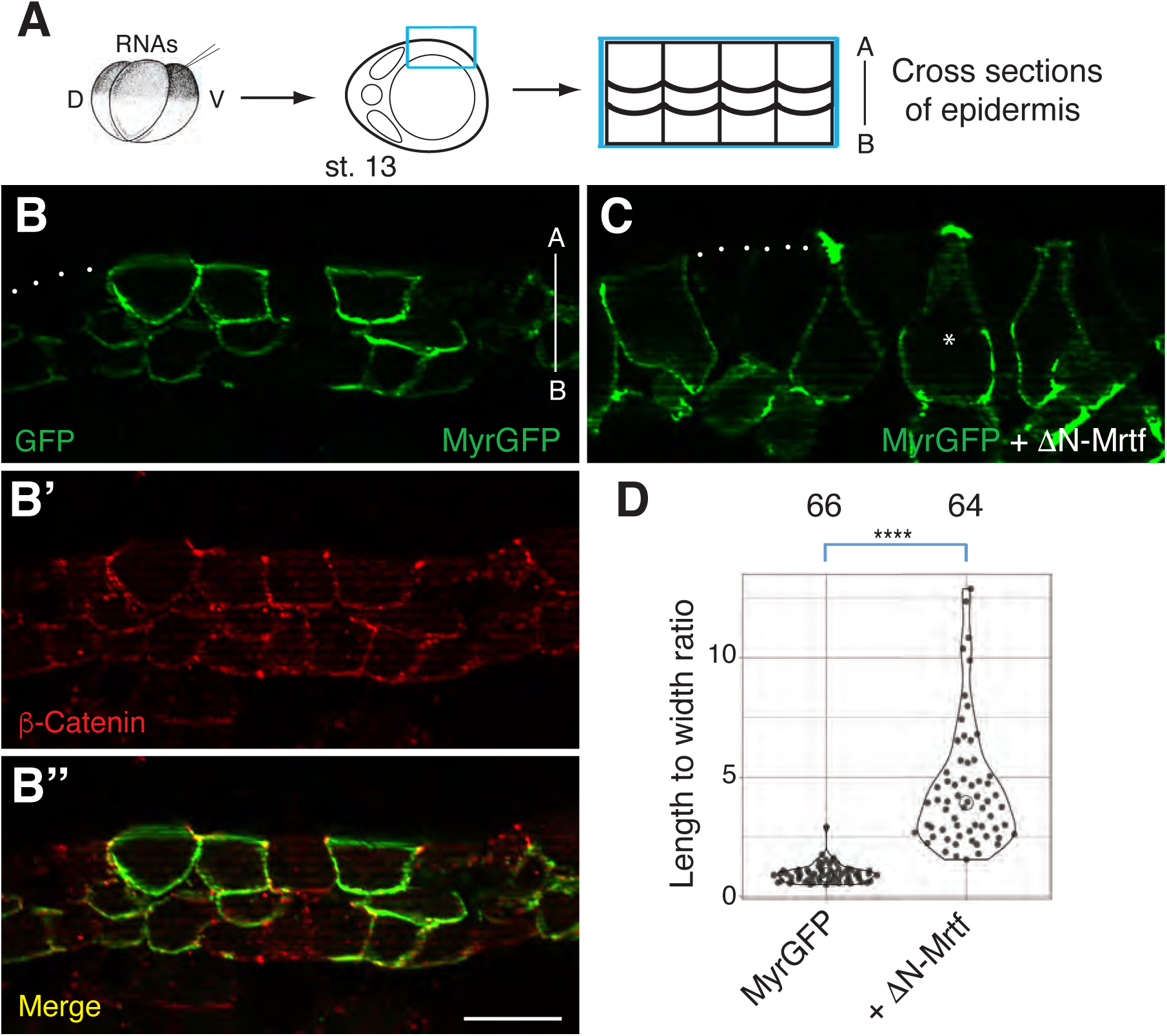
Bottle cell morphology of ΔN-Mrtf expressing cells. A. Experimental scheme. Embryos were injected with 250 pg of ΔN-Mrtf RNA and 50 pg of MyrGFP RNA as indicated in Fig. 5 legend. MyrGFP is a lineage tracer. B, C, Transverse cryosections of control (B) or ΔN-Mrtf-expressing (C) ventral ectoderm (stage 13) immunostained for GFP and β-catenin. Punctate lines indicate apical cell surface. Scale bar: 50 μm. D. Quantification of the ratio between cell length along the apical-basal axis and apical domain width. Numbers of scored cells are indicated on top. ****p<0.0001, Student’s t-test.

These results indicate that active Mrtf promotes ectopic apical constriction during gastrulation. Notably, ΔN-Mrtf did not cause hyperpigmentation that marks the effect of other apical constriction inducers, including Plekhg5, GEF-H1 or LMO7 (Itoh et al., 2014; Matsuda et al., 2022; Popov et al., 2018). This suggests that a distinct molecular mechanism may be involved.

To gain additional insights, we evaluated whether the induction of apical constriction by ΔN-Mrtf requires it transcriptional activity. We therefore tested ΔNΔC-Mrtf that lacks transcription activation domain and is expressed at similar levels in ectoderm cells (**Fig. S3**). ΔNΔC-Mrtf was unable to trigger apical constriction (**Fig. 5E-E’’, H**). In another experiment, ΔN-Mrtf activity was completely inhibited by the coexpression of ΔC-Mrtf (**Fig. S5**). These results indicate that transcriptional mechanisms are crucial for the induction of apical constriction by Mrtf.

We also tested whether Mrtf would regulate cell intercalations in ectoderm explants after stimulation with Activin, a Nodal/TGF beta superfamily signaling factor. Ligand-stimulated activin receptors signal by phosphorylating Smad2/Smad3 and can lead to ectodermal explant elongation that is thought to mimic convergent extension movements (Howard and Smith, 1993; Sokol, 1996). We wanted to know whether Mrtf constructs influence Activin-dependent elongation of ectoderm explants (**Fig. S6A-H**). All Mrtf constructs significantly inhibited explant elongation and ΔN-Mrtf completely blocked it (**Fig. S6E-H**). Notably, we did not see changes in Smad2 phosphorylation (**Fig. S6I).** This confirms that ΔN-Mrtf affects cell behavior rather than the ability of Activin to signal to its receptors. Together, our observations show that, while ΔN-Mrtf triggers ectopic apical constriction, it inhibits elongation of Activin-treated ectoderm explants, pointing to potential roles of Mrtf in additional cell behaviors.

## DISCUSSION

Our study addressed a role of Mrtf signaling in the control of morphogenetic processes during *Xenopus* gastrulation and neurulation. We observed abnormal blastopore formation in embryos depleted of *Mrtfa* and *Mrtfb* and those expressing dominant interfering Mrtf constructs. At later stages, the embryos with reduced Mrtf activity exhibited neural tube abnormalities. These defects have not been described in mice lacking *mrtfa* or *mrtfb* genes, likely due to the functional redundancy. We note that dorsal blastopore was delayed but not fully disrupted by Mrtf knockdown, possibly because Mrtfa and Mrtfb are maternally expressed (www.Xenbase.org).

We sought to understand how Mrtfs regulate morphogenesis. Although embryos with decreased and elevated Mrtf activity exhibited similar morphological abnormalities, the effects of the oppositely acting Mrtf constructs on F-actin distribution and cell behavior were different. Suppressing Mrtf-dependent transcription reduced overall F-actin levels and inhibited epithelial folding during gastrulation and neurulation, suggesting an apical constriction defect. By contrast, a constitutively active Mrtf inhibited F-actin at tricellular junctions (TCJ), likely leading to their remodeling. At early neurula stages, we observed that the cells acquired convex shape, reduced their apical surface and initiated apical constriction.

We hypothesize that Mrtf controls morphogenesis by reorganizing F-actin at TCJ Consistent with this possibility, Mrtf gene targets include actomyosin modulators and cell junction molecules (data not shown). TCJ remodeling is thought to be responsible for enhanced actomyosin contractility and has been previously associated with rounded cell shape due to increased cell tension and apical domain reduction (Arnold et al., 2019; Bosveld and Bellaiche, 2020; Cho et al., 2022; Higashi and Miller, 2017). One would expect an enhanced localization of actomyosin complexes at TCJ in apically constricting cells. Surprisingly, F-actin and Tricellulin localization at TCJ have been decreased in response to ΔN-Mrtf, suggesting a different, yet uncharacterized mechanism. We hypothesize that stimulation of G-actin transcription by Mrtf may promote vesicular trafficking or another actin-dependent process leading to TCJ disassembly and apical domain reduction. In addition, it is possible that, besides actins, other Mrtf gene targets identified by our RNA sequencing analysis are involved in the apical constriction phenotype.

The observed apical domain reduction in response to ΔN-Mrtf RNA injection is reminiscent of enhanced contractility of MDCK cells lacking the ZO1 protein (Choi et al., 2016; Fanning et al., 2012). Of note, we observed that apical constriction triggered by ΔN-Mrtf is not accompanied by hyperpigmentation, a common feature that accompanies membrane shrinking in response to several apical constriction inducers, such as Plekhg5, Gef-H1 or Lmo7 (Itoh et al., 2014; Matsuda et al., 2022; Popov et al., 2018). Our unexpected findings highlight mechanistic differences between cell intercalations and apical constriction. Whereas both cell behaviors involve actomyosin contractility, Mrtf promotes apical constriction but inhibits the elongation of Activin-treated explants. This activity is consistent with the delay in blastopore closure in overexpressing embryos. The observed inhibition of cell intercalations may be due to inability of the constricting cells to intercalate, or could be caused by yet uncharacterized activity of Mrtf. Future studies are warranted for better understanding of Mrtf functions in various developmental processes.

## Materials and Methods

### Plasmids, in vitro RNA synthesis and morpholino oligonucleotides

pCS2-Flag-Mrtf plasmids have been generated by PCR from the *X. laevis* DNA clone for *mrtfa.S* (accession number NM_001088851) obtained from Horizon. ΔN-Mrtf (lacking aa6-211), ΔB-Mrtf (lacking aa323-367), ΔC-Mrtf (lacking aa594-991) were produced using single primer mutagenesis with specific primers (Makarova et al., 2000). ΔNΔC-Mrtf was generated from ΔN-Mrtf. Details of cloning are available upon request. All constructs have been confirmed by sequencing.

Capped mRNAs were synthesized using the Ambion mMessage mMachine kit (ThermoFisher). Linearized RNA templates were made from pCS2-FlagMrtf derivatives, pCS2-memRFP, pCS2-LifeactGFP, pCS2-myr-tagGFP, pCS2-myr-tagRFP, pCS2-Tricellulin-mCherry and pCS2-FlagGFP.

Splicing-blocking morpholinos (MOs), Mrtfa MO^sp^ and Mrtfb-MO^sp^, were purchased from Gene Tools (Philomath, OR). The MOs had the following sequences: Mrtfa MO^sp^, 5’-GGTGACTGGGACCTGAAACAGAAAT-3’; Mrtfb-MO^sp^, 5’-CAGCCGGACTCTTTAATGCTGAAAG-3’.

### Xenopus embryo culture, microinjections and animal cap assay

Wild-type *Xenopus laevis* were maintained following the Guide for the Care and Use of Laboratory Animals of the National Institutes of Health. A protocol for animal use was approved by the Institutional Animal Care and Use Committee (IACUC) of the Icahn School of Medicine at Mount Sinai. *In vitro* fertilization and culture of *Xenopus laevis* embryos were carried out as previously described (Itoh et al., 2005). Staging was according to (Nieuwkoop and Faber, 1967). For microinjections, 2 to 8-cell embryos were transferred into 3 % Ficoll in 0.5 x Marc’s Modified Ringer’s (MMR) solution (50 mM NaCl, 1 mM KCl, 1 mM CaCl_2_, 0.5 mM MgCl_2_, 2.5 mM HEPES pH 7.4) (Peng, 1991) and 10 nl of *3DA-Luc* DNA (a gift of G. Posern), mRNA or MO solution were injected into one, two or four blastomeres of 2-8-cell embryos, either animally or sub-equatorially. Injected embryos were transferred into 0.1 x MMR at blastula stages. Amounts of injected mRNA per embryo have been optimized in preliminary dose-response experiments and are indicated in figure legends.

For ectodermal explant assay, 4-8 cell embryos were injected with RNAs into four animal blastomeres and ectodermal explants were excised at stage 8-8.5. The explants were incubated in the presence or absence of 0.5 ng/ml of activin in low Ca/Mg MMR solution (Itoh and Sokol, 1994) with 0.1 mg/ml BSA for 1 hr and processed for immunoblotting or cultured in 0.5 x MMR until control embryos reached stage 15 to observe explant elongation phenotypes.

### RNA sequencing and RT-PCR

For RT-PCR, two cell embryos were injected with Mrtfa-MO^sp^ or Mrtfb-MO^sp^ and cultured until stage 15. RNA was extracted from 5 embryos, using the RNeasy kit (Qiagen). For RT-PCR, cDNA was made from 1-2 µg of total RNA using the first strand cDNA synthesis kit (Invitrogen) according to the manufacturer’s instructions. Primers used for RT-PCR are listed in **Suppl. Table 1**.

For RNA sequencing, Flag-ΔN-Mrtf (0.5 ng) RNA was injected twice into animal pole region of two cell embryos. Ectoderm explants were prepared from injected and control uninjected embryos when siblings reache stage 9-9.5 and cultured until stage 11. RNA was extracted from 30 ectoderm explants at stage 11 using RNeasy kit (Qiagen). cDNA library preparation, paired-end 150 bp sequencing using Illumina HiSeq2000 analyzers and bioinformatics analysis were performed by Novogene (Sacramento, CA). The raw reads were filtered to remove reads containing adapters and of low quality. RNA counts were normalized to an uninjected control set. Sequences were mapped to the *Xenopus* genome version XL-9.1_v1.8.3.2 at http://www.xenbase.org/other/static/ftpDatafiles.jsp using Hisat2. The differentially expressed genes (DEGs) were detected using DESeq (Anders and Huber, 2010) with the two-fold change cutoff. The p-value estimation was based on the negative binomial distribution, using the Benjamini-Hochberg estimation model with the adjusted p < 0.05. RNA sequencing data have been obtained in three independent experiments. A representative dataset has been submitted to NCBI GEO (accession GSE243351).

For RT-qPCR, two-cell embryos were injected animally with 500 pg ΔN-Mrtf RNA and cultured until stage 9-9.5 when ectodermal explants were excised. When sibling embryos reached stage 11, the explants were harvested for total RNA extraction. cDNA was made as described above and the reactions were amplified using a CFX96 light cycler (Bio-Rad) with Universal SYBR Green Supermix (Bio-Rad). Primer sequences used for RT-qPCR are shown in **Suppl. Table 1**. The reaction mixture consisted of 1X Power SYBR Green PCR Master Mix, 0.3 μM primers, and 1 μl of cDNA in a total volume of 10 μl. The ΔΔCT method was used to quantify the results. All samples were normalized to control explants. Transcripts for *elongation factor 1a1 (ef1a1*) were used for normalization. Data are representative of two to three independent experiments and shown as means +/- s.d.

### Fluorescent imaging and phalloidin staining

For apical cell domain size analyses, 4-8 cell embryos were injected with 50 pg Myr-GFP or Myr-RFP and 250 pg ΔN-Mrtf, 1 ng ΔC-Mrtf or ΔNΔC-Mrtf RNAs into one ventro-animal blastomere. The embryos were fixed in 3.7% formaldehyde in PBS for 30 min when control embryos reached indicated stages. Ventral ectoderm was excised and imaged. To detect F-actin, 4-8 cell embryos were injected with RNAs into one dorso-vegetal blastomere and the injected embryos were fixed in 3.7% formaldehyde in PBS for 30 min when control embryos reached stage 10. After washing with PBS, dorsal marginal zone (DMZ) explants were excised and incubated with Alexa488-phalloidin or Alexa-568-phalloidin in PBS-Tween overnight at 4°C. Fluorescence imaging was carried out using AxioImager (Zeiss) microscope with the Apotome attachment. Alternatively, ectodermal explants were imaged with the LSM880 confocal microscope (Zeiss) using MSSM Core facility.

### Time lapse imaging of Xenopus embryos

For time-lapse imaging, 4-8 cell embryos were injected with 250 pg ΔN-Mrtf or ΔC-Mrtf RNA with 50 pg Myr-GFP or 100 pg Lifeact-GFP RNA into one ventro-animal blastomere and cultured until stage 10.25 or 11. The injected embryos were mounted in 1% low melting agarose (Lonza) on a slide coverslip attached to a silicone isolator (Grace Bio-labs) or on a glass-bottom dish (Cellvis). Time-lapse imaging was carried out using the AxioZoom V16 fluorescence stereomicroscope (Zeiss) equipped with the AxioCam 506 mono CCD camera (Zeiss) or the LSM880 confocal microscope (Zeiss). Images were taken every 5 or 8.5 min over the period of 1.5 or 3.5 hrs. Apically constricting cells were defined as the ones in which the apical domain size has been reduced by more than 20%.

### Cryosectioning and immunostaining

For cryo-sectioning, embryos were devitellinized at stage 13, fixed in 3.7% formaldehyde/PBS or Dent’s fixative for 1-2 hrs and embedded in 15% fish gelatin/15% sucrose solution (Itoh et al., 2014). The embedded embryos were frozen in dry ice and sectioned at 10 μm using Leica CM3050 cryostat. Cryo-sections were stained with anti-GFP (Santa Cruz, 1/200) and anti-β-Catenin (Sigma, 1/200) antibodies and Alexa488-conjugated (Invitrogen, 1/200) or Cy3-conjugated (Jackson Immunoresearch, 1/200) secondary antibodies. Fluorescence of the stained cryosections was examined under AxioImager (Zeiss) microscope with the Apotome attachment.

### Quantification and statistical analyses

To quantify apical domain size, explants expressing membrane marker MyrGFP, MyrRFP or or LifeactGFP were imaged. After image acquisition, the apical domains of individual cells were manually outlined using a free-hand line tool in ImageJ software based on membrane fluorescence. Only mosaically expressing cells with less than three contacts with other expressing cells were used for quantification. To quantify phalloidin intensities of the tricellular junctions, bicellular junctions or cytoplasm, explants expressing memRFP were stained with Phalloidin-488. After image acquisition, a square tool was used in ImageJ to measure Phalloidin intensities at different cellular locations.

To quantify convex area, live imaging of embryos expressing LifeactGFP in the superficial layer of ectoderm at stage 11 was used. Convex membrane curvature area was manually outlined between two junctional spots of Lifeact-GFP using a free-hand line tool and scored as positive value. Concave membrane curvature area was similarly outlined but scored as negative value.

To quantify length-to-width ratios in animal cap explants, images of animal cap elongation in the presence or absence of Activin were acquired when sibling embryos reached stage 15. Length along animal cap tissue elongation and cap width were measured using a free-hand line tool in ImageJ. To quantify length-to-width ratios of superficial ectoderm in cross-sections, cell length along apical to basal axis and cell width at the apical surface were measured using the same tool in Image J.

For statistical analysis, Student’s t-test was used to determine statistical significance between two groups. Non-significant, n.s. (p>0.05). *p<0.05, **p<0.01, ***p<0.001, ****p<0.0001.

### Luciferase activity assays and immunoblotting

For luciferase activity assays, 20 pg of *3DA-Luc* reporter plasmid DNA (Posern et al., 2002) was coinjected with RNAs into two animal blastomeres of 4-8 cell embryos. Luciferase activity was measured when sibling embryos reached stage 10+ to 10.5 as described (Itoh et al., 2014).

For immunoblot analysis, 4-8-cell embryos were injected with RNAs encoding Mrtf constructs into two or four animal blastomeres. Whole cell lysates were prepared from five embryos with 85 μl of the lysis buffer (50 mM Tris-HCl pH 7.6, 50 mM NaCl, 1 mM EDTA, 1% Triton X-100, 10 mM NaF, 1 mM Na_3_VO_4_, 1 mM PMSF) when sibling embryos reached stage 10.5 and the lysates were collected after centrifugation for 4 min at 16,000 g. For immunoblot analysis of animal caps, five animal caps were lysed with 25 μl of the lysis buffer. Protein lysates were separated by SDS-PAGE and subjected to immunoblotting as previously described (Itoh et al., 2005). The following primary antibodies were used: mouse anti-FLAG (M2, Sigma), rabbit anti-Erk1/2 (9102, Cell Signaling), rabbit anti-phospho-Smad2 (3108S, Cell Signaling) or rabbit anti-β-catenin (71-2700, Invitrogen) antibodies. The detection was carried out by enhanced chemiluminescence as described (Itoh et al., 2005), using the ChemiDoc MP imager (BioRad).

## Acknowledgements

We thank Ron Prywes for the human Mkl-1 plasmids used in initial experiments, Guido Posern and Ann Miller for plasmids, Mark Corkins for advice on RNA sequence analysis and members of the Sokol laboratory for discussions. We acknowledge the help from the ISMMS Microscopy Core facility. This study has been supported by the NIH grants R01DE027665 and R35GM122492 to SYS.

## Supplementary figures S1-S6

**Fig. S1.**
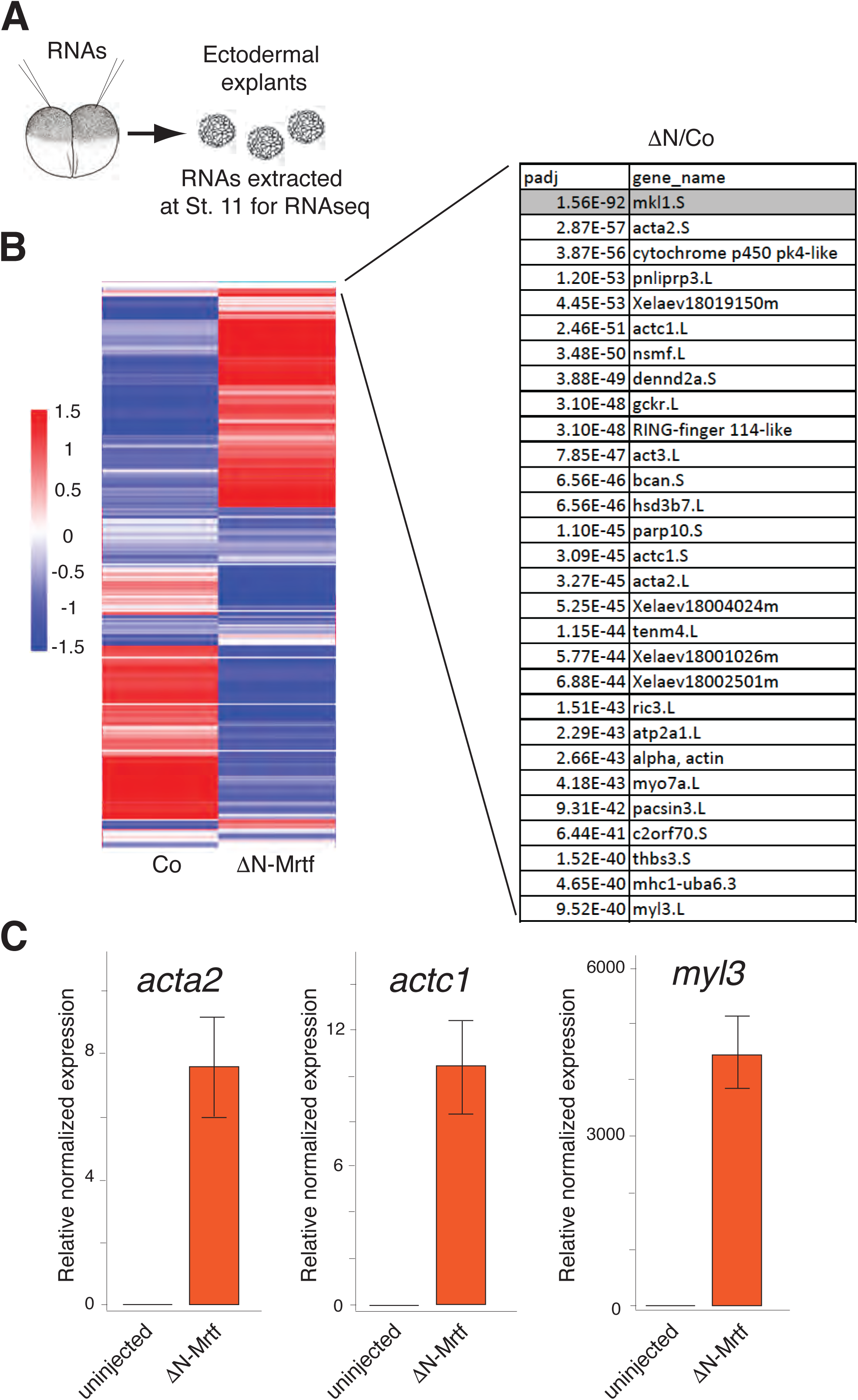
Transcriptome analysis identifies putative Mrtf target genes. A, Experimental scheme. Two-cell embryos were injected into the animal pole region with 500 pg of ΔN-Mrtf RNA. Ectodermal explants were excised at stage 9-9.5. Total RNA was extracted when sibling embryos reached stage 11 for sequencing. B, Heatmaps show relative expression of differentially expressed genes in the control and ΔN-Mrtf expressing ectoderm. Top enriched genes are shown on the right, with adjusted p-value (Padj) indicated. C. Validation of *acta2, actc1* and *myl3* regulation by ΔN-Mrtf using quantitative RT-PCR.

**Fig. S2.**
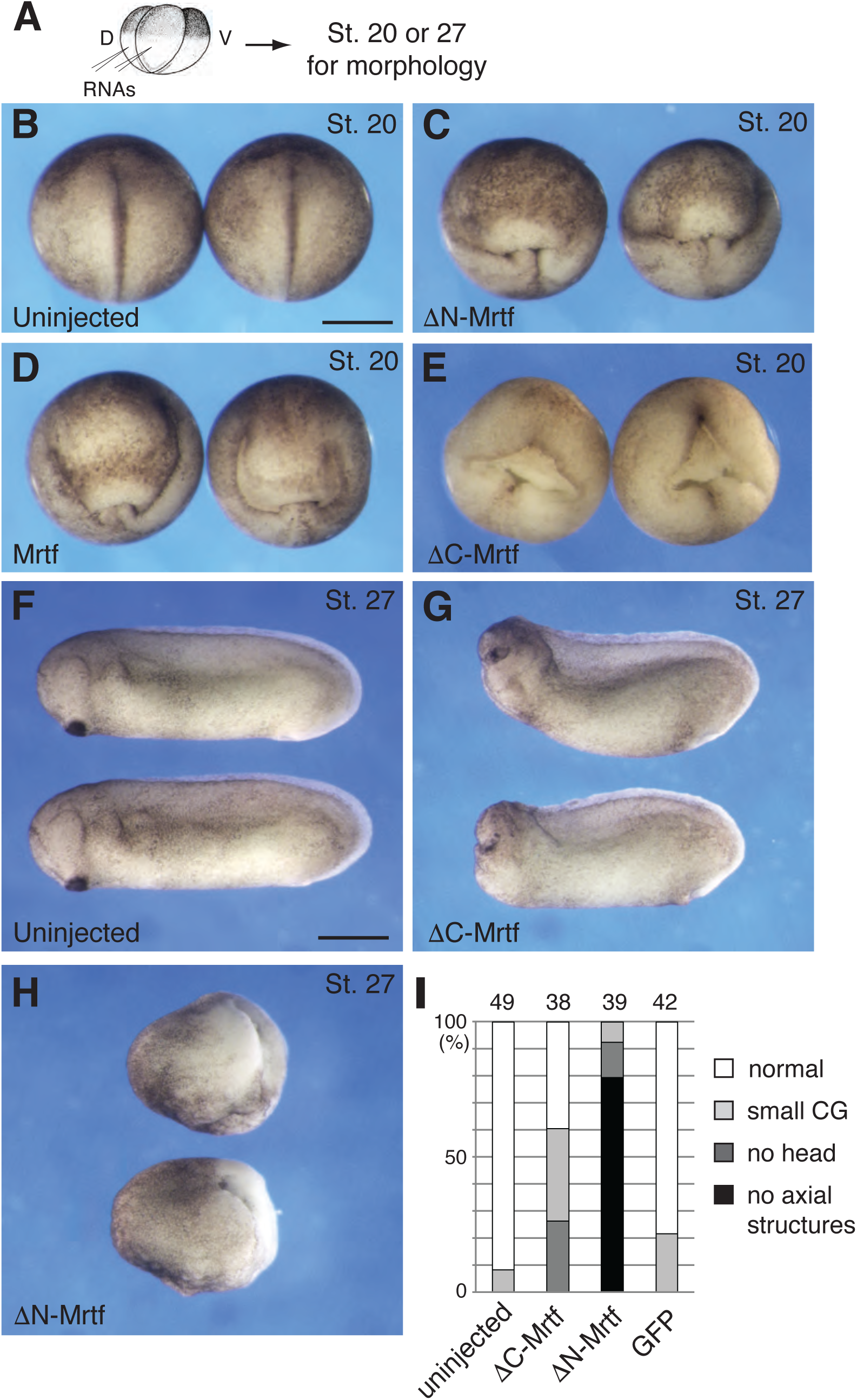
Neurulation defects caused by active and dominant interfering Mrtf constructs. A. Experimental scheme. Four-cell embryos were left untreated (B) or injected dorso-vegetally with 166 pg ΔN-Mrtf RNA (C), 500 pg or 1 ng Mrtf RNA (D), or 1 ng ΔC-Mrtf RNA (E). Representative images were taken when control embryos reached stage 20 (B-E, dorsal view) or stage 27 (F-H, lateral view). Scale bar (in B, F, applies everywhere): 500 μm. I, Quantification of the developmental abnormalities for stage 27 embryos injected with Mrtf RNAs. Dorsoanterior defects were scored as microcephalic (small cement gland), headless, or lacking anterior and dorsal axial structures. Data are representative of three independent experiments.

**Fig. S3.**
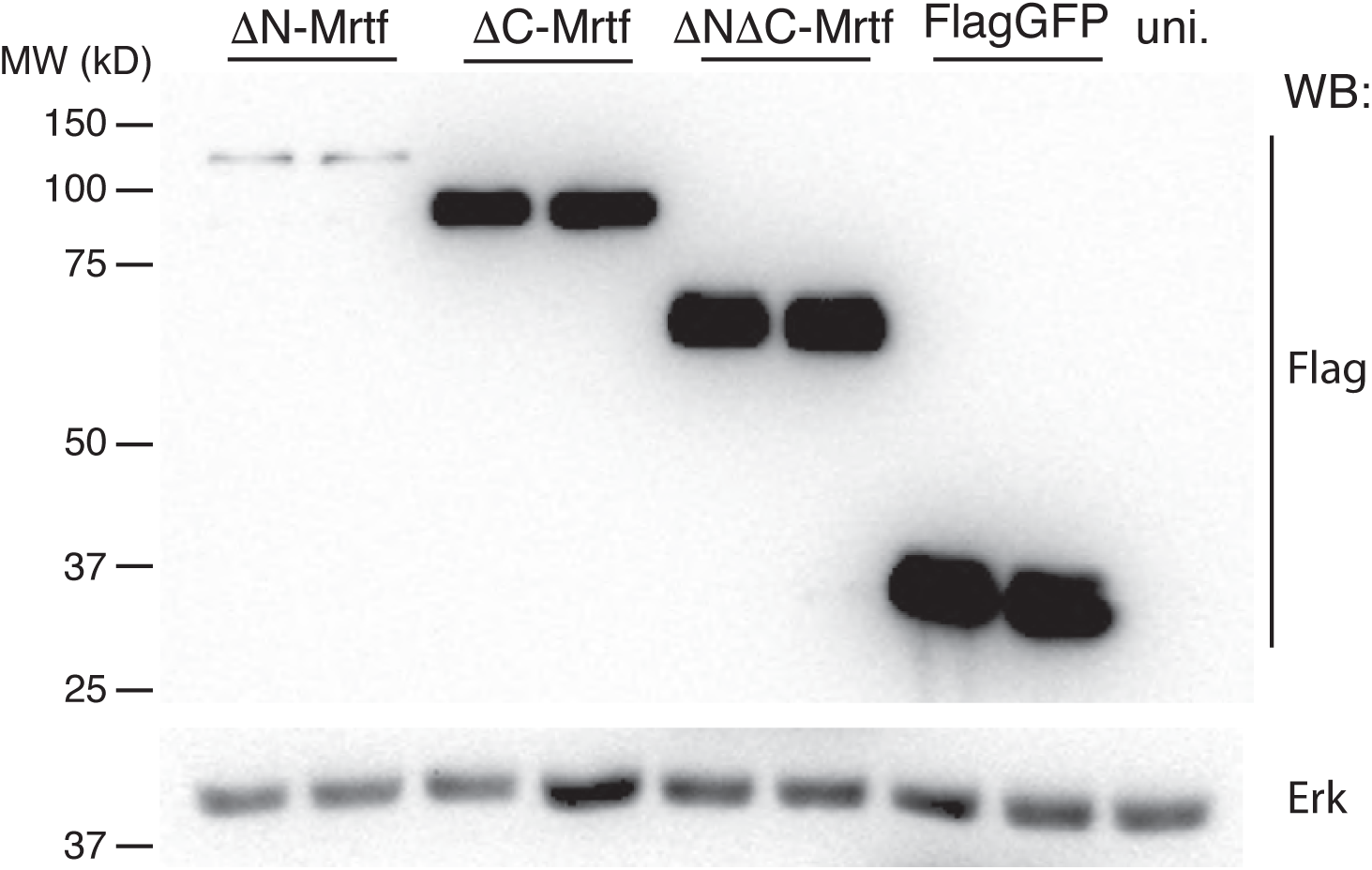
Expression levels of Mrtf constructs in *Xenopus* embryos. Four animal blastomeres at 4-8 cell stage were injected with the following RNAs as indicated, ΔN-Mrtf (500 pg), ΔC-Mrtf (1 ng) ΔNΔC-Mrtf (1 ng) or FlagGFP (1 ng). When control embryos reached stage 10.5, injected embryos were lysed for immunoblotting with anti-Flag or anti-Erk1 antibodies. Erk1 serves as a loading control.

**Fig. S4.**
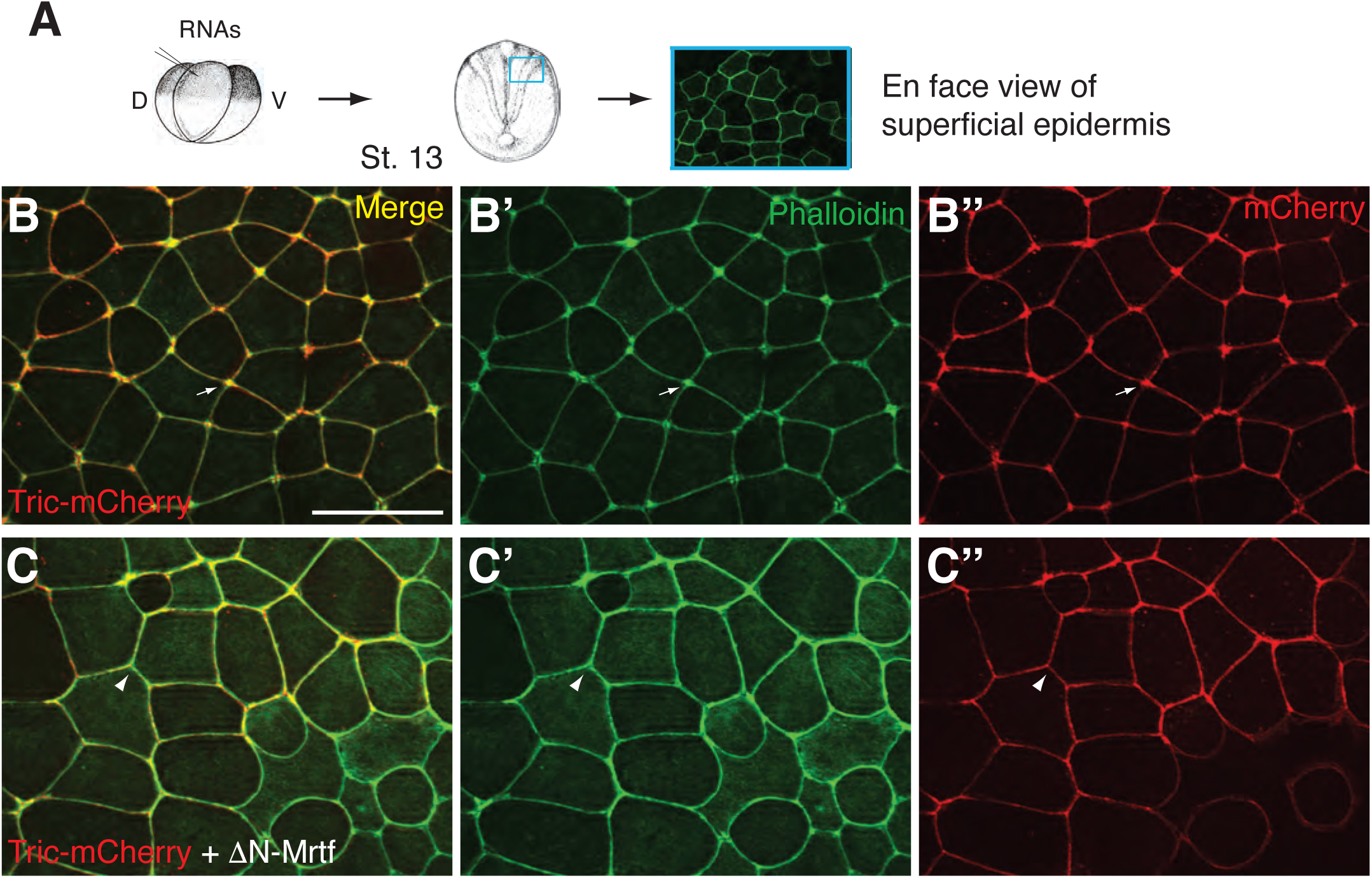
Mrtf-induced apical constriction involves tricellular junction remodeling. A. Experimental scheme. Four-cell embryos were injected into one dorso-animal blastomere with Tricellulin (Tric)-mCherry RNA (25 pg) alone (B-B”) or with 250 pg ΔN-Mrtf RNA (C-C”). Superficial cells are imaged for phalloidin staining (B’, C’) or mCherry expression (B’’, C’’). Arrows in B-B” indicate TCJ. Arrowheads in C-C” indicate reduction of F-actin and Tricellulin at TCJ. Note the spreading of the Tric-mCherry and F-actin signals to the bicellular junctions in B, C.

**Fig. S5.**
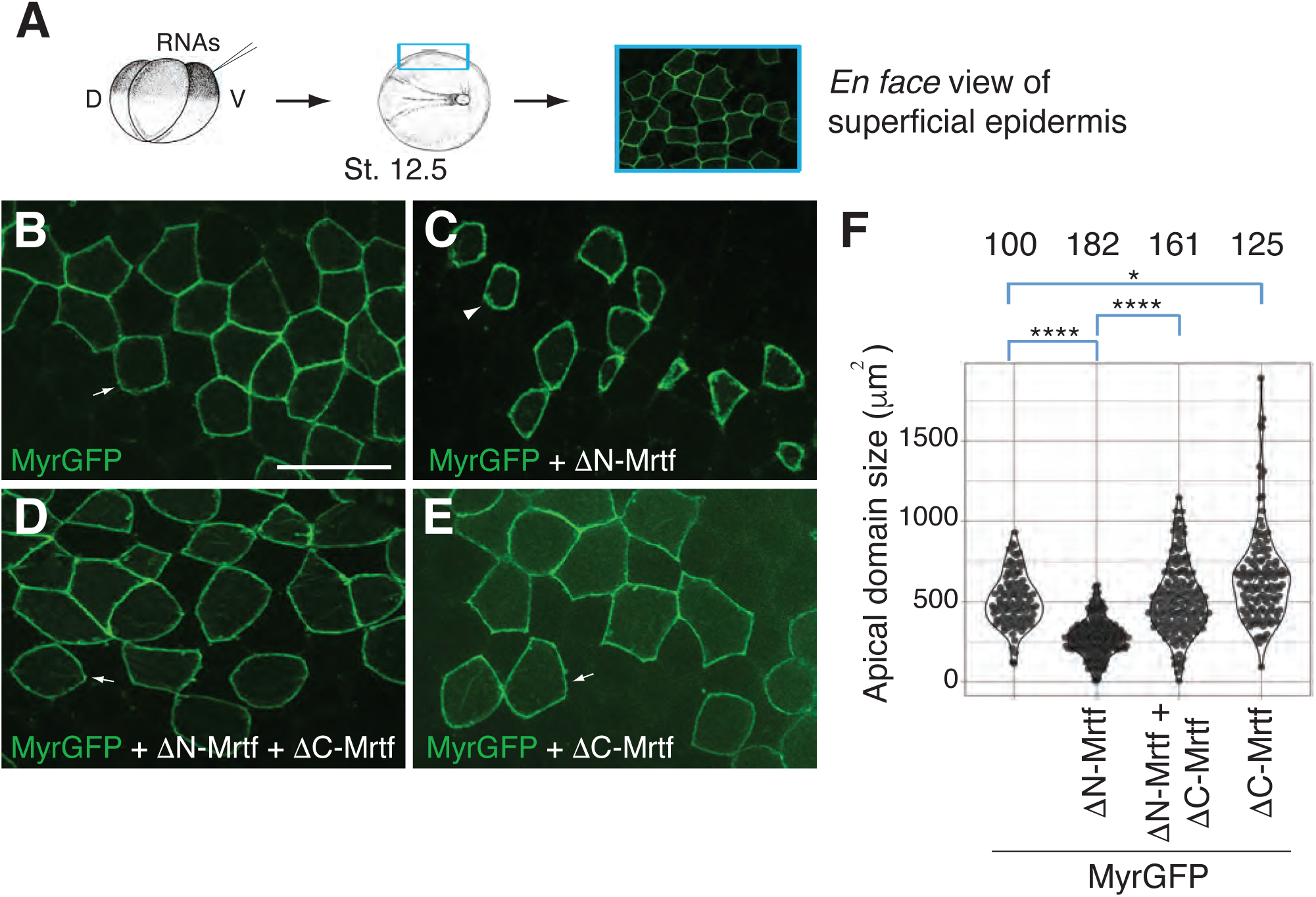
Mrtf lacking transcriptional activation domain counteracts Mrtf-induced apical constriction. A. Experimental scheme. Four-cell embryos were injected into one-ventro-animal blastomere with MyrGFP RNA (50 pg) alone (B) or with 250 pg ΔN-Mrtf RNA (C), with 250 pg ΔN-Mrtf and 1 ng ΔC-Mrtf RNAs (D) or with 1 ng ΔC-Mrtf RNA (E). Superficial epidermal ectoderm of the injected embryos at stage 12.5 was imaged. F. Apical domain size of mosaic GFP expressing cells in B-E was measured. Arrows (B, D, E) indicate mosaic cells. Arrowhead in C indicates an example of reduced size of a mosaic cell. Numbers of scored cells are on the top. *p<0.05, ****p<0.0001, Student’s t-test. Scale bar: 50 μm.

**Fig. S6.**
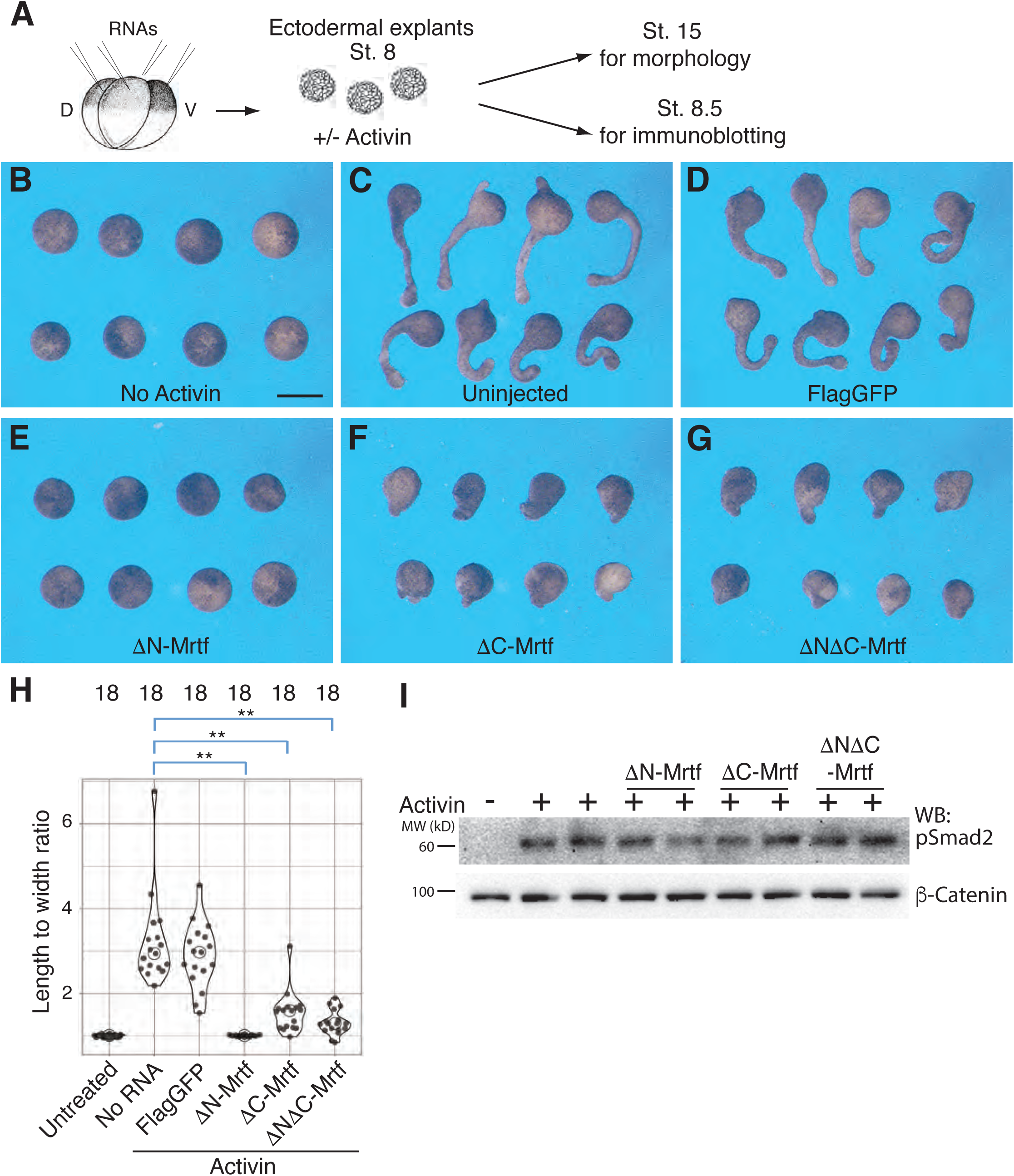
Interference with Mrtf activity inhibits activin-dependent elongation of ectodermal explants. A, Experimental scheme. Four animal blastomeres of 4-8 cell embryos were injected with FlagGFP (1 ng), ΔN-Mrtf (500 pg), ΔC-Mrtf (1 ng) or ΔNΔC-Mrtf (1 ng) RNA. When control embryos reached stage 8, ectodermal explants were excised and treated with activin. The explants were cultured until control embryos reached stage 15 for observation of animal cap morphology (B-G) or collected at stage 8.5 for immunoblot analysis (I). Control untreated explants (B), activin-treated explants (C-G). Typical effects of Mrtf constructs on ectodermal explants are shown. (H) Quantification of length to width ratio of the explants shown in B-G and in data not shown. Numbers of scored explants are indicated on the top. I, Activin-dependent increase in Smad2 phosphorylation is unaffected by Mrtf manipulation. Immunoblot was probed with anti-phospho-Smad2 or anti-β-catenin antibodies as a control for loading. The data are representatives of 3-5 independent experiments. Scale bar: 300 μm.

**Supplementary Table 1.**
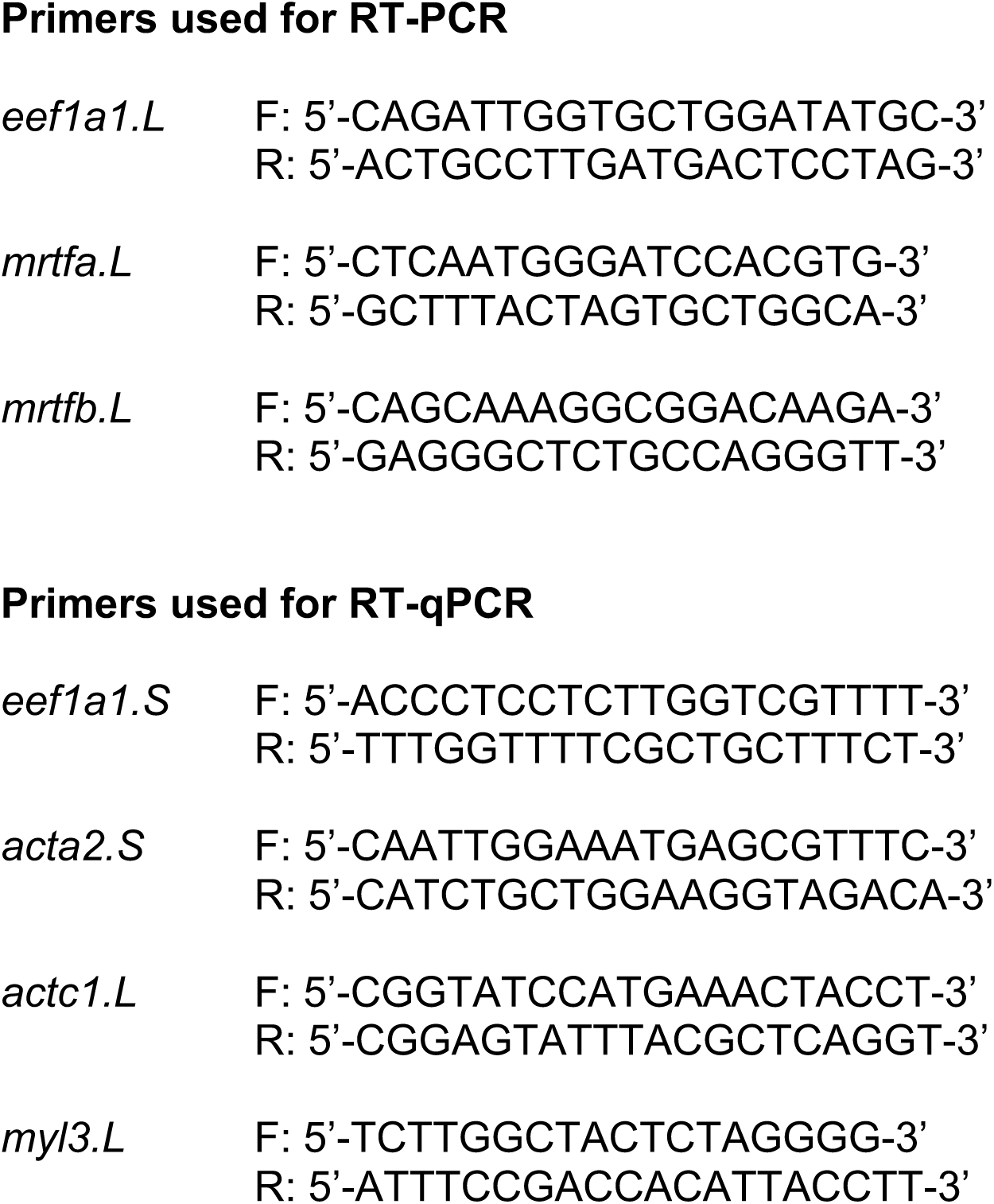
List of RT-PCR primers used in this study.

## Supplementary movies

**Supplementary Movie 1.**

Time-lapse live imaging of superficial nonneural ectoderm cells expressing LifeAct-GFP RNA from stage 11 embryos using the Zeiss LSM880 confocal microscope (20 x magnification lens). Length of the movie 80 min.

**Supplementary Movie 2.**

Time-lapse live imaging of nonneural ectoderm cells expressing LifeAct-GFP and ΔN-Mrtf RNA from stage 11 embryos using Zeiss LSM880 confocal microscope (20 x magnification lens). Length of the movie 80 min.

